# Evolution-Guided Engineering of Non-conserved *N*-Glycosylation Sites Improves Heterologous Production of Sulfotransferases

**DOI:** 10.64898/2026.10.18.759982

**Authors:** Chao-Yue Li, Yi Li, Xin-Yu Li, Yu-Han Zhao, Jiu-Ying Sun, Ya-Lin Cao, Xue-Ping Guo, Shuang Zheng, Jian-Qun Deng, Ju-Zheng Sheng

## Abstract

*Escherichia coli* features low cost, mature fermentation technology and high recombinant protein output, but the expression of glycosylation-dependent eukaryotic proteins therein is severely limited. Many glycosylated proteins are key tool enzymes for the biosynthesis of various pharmaceutical molecules, such as glycosaminoglycan-related synthases, but there is a lack of universal protein engineering methods to enable the adaptive evolution of these enzymes in *E. coli*. We established an evolution-guided optimization strategy that targets non-conserved *N*-glycosylation sites (EGO-*N*Gly). The core of this strategy lies in the phylogeny-guided selective reversion of evolutionarily acquired *N*-glycosylation sequons to naturally occurring non-glycosylated residues from early-divergent orthologs, thereby improving heterologous production and residual activity stability for selected sulfotransferases in *E. coli*. Using heparin 3-*O*-sulfotransferase 1 (3OST-1) as a model enzyme apparatus in eukaryotic organisms, EGO-*N*Gly extended the residual activity half-life by 4.5-fold. Structural analysis and molecular dynamics simulations indicate that greater surface rigidity and remodeling of intramolecular interaction networks contribute to the improved conformational stability of M7 in an aglycosylated host. EGO-*N*Gly was further applied to a panel of 3OST and 6OST homologs, demonstrating its applicability to a subset of the tested enzymes. Furthermore, EGO-*N*Gly can be orthogonally integrated with other complementary protein engineering strategies, as exemplified by its combination with the machine learning-aided Protein Repair One-Stop Service, generating variants with 8-fold and 17-fold extensions in residual activity half-life. Collectively, these results establish EGO-*N*Gly as a practical and potentially cost-effective engineering strategy for improving the prokaryotic expression and functional performance of selected glycosylation-dependent eukaryotic sulfotransferases.

**Significance Statement:** *Escherichia coli* lacks rational engineering methods for the heterologous expression of glycosylation-dependent eukaryotic proteins. We developed an evolution-guided optimization of *N*-glycosylation (EGO-*N*Gly) strategy that reverts non-conserved glycosylation sequons to the non-glycosylated residues conserved in early-divergent orthologs, improving the heterologous expression, catalytic function, and stability of animal-derived sulfotransferases. In heparin 3-*O*-sulfotransferase 1, EGO-*N*Gly extended the residual activity half-life at 37 °C by 4.5-fold. The strategy was transferable to several additional sulfotransferase homologs in a sequence-context-dependent manner and could be combined with machine-learning aided stability design to further extend the half-life by up to 17-fold. EGO-*N*Gly thus provides a practical and potentially cost-effective strategy for improving the prokaryotic production of selected animal-derived sulfotransferases.

## Introduction

*Escherichia coli* is the most widely used heterologous host for industrial enzyme production, owing to its low cost, mature high-density fermentation technology, and high recombinant protein yield (1–3). However, *E. coli* lacks the canonical *N*-linked glycosylation system that is ubiquitous in eukaryotes (4, 5). As a result, many eukaryotic proteins containing *N*-glycosylation sites are prone to misfolding, low soluble expression, or loss of activity in this host (6–9) (Figure 1a). Notably, eukaryotic sulfotransferases typically contain multiple *N*-glycosylation sequons, and their folding and functional stability often depend on *N*-glycosylation. Thus, glycosylation-dependent expression remains a major bottleneck for prokaryotic production of eukaryotic enzymes.

**Figure 1.**
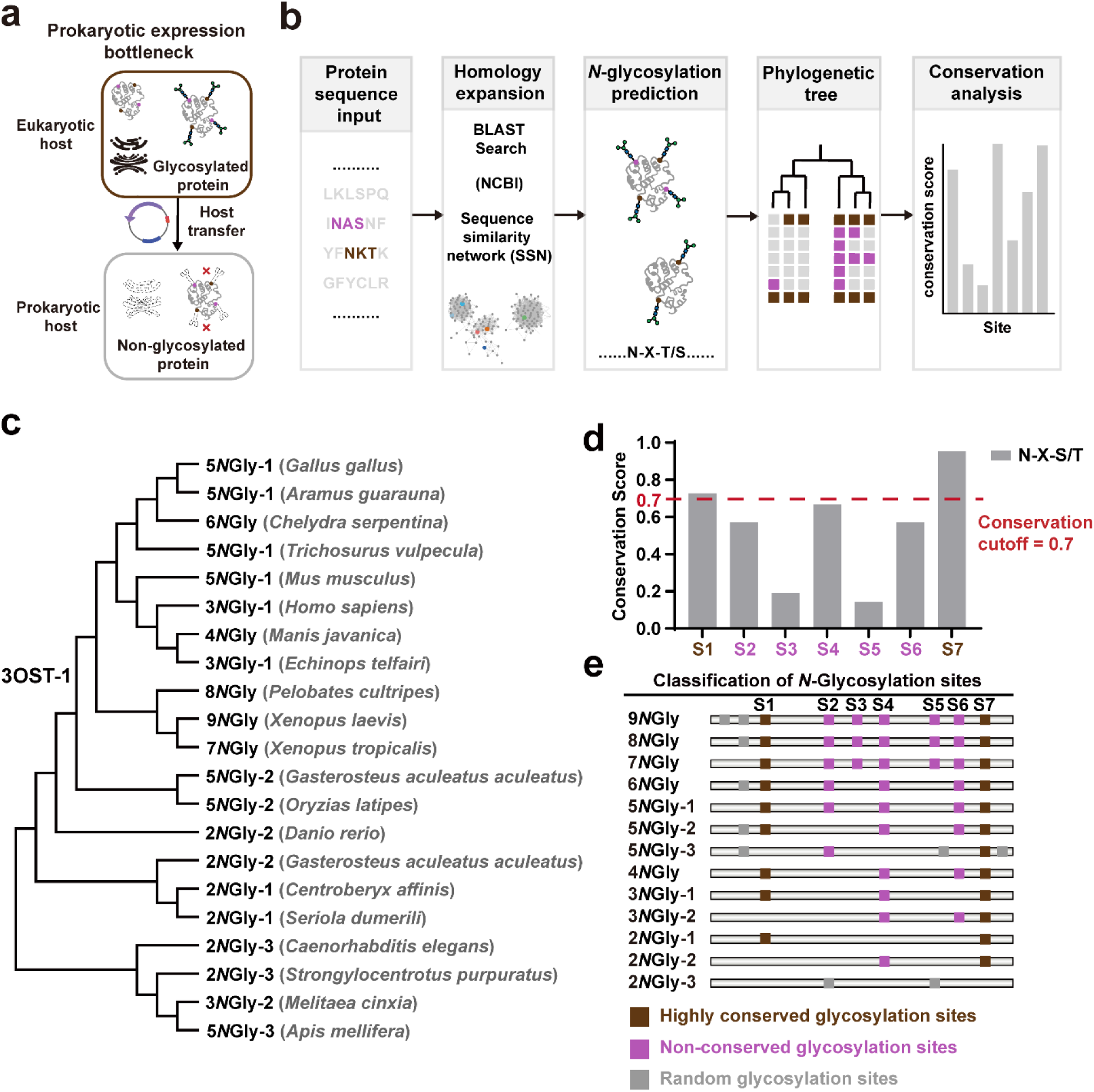
Prokaryotic expression bottleneck and evolutionary analysis of 3OST-1 *N*-glycosylation sites. (a) Schematic illustration of the prokaryotic expression bottleneck for eukaryotic glycoproteins. (b) Identification workflow for candidate *N*-glycosylation sequons in 3OST-1. Sequence analysis integrating BLAST, sequence similarity network (SSN), *N*-glycosylation prediction, phylogenetic reconstruction, and conservation analysis identified seven potential *N*-glycosylation sequons in 3-*O*-sulfotransferase 1 (3OST-1) (c) Phylogenetic distribution of 3OST-1 *N*-glycosylation sites across different evolutionary lineages. (d) Conservation scores of the seven candidate *N*-glycosylation sites. For each site, the conservation score represents the frequency of the canonical N-X-S/T sequon (X ≠ Pro) at the corresponding aligned position in the ortholog multiple-sequence alignment. The red dashed line indicates the operational conservation cutoff (0.7) used in this study for candidate stratification. Sites with scores ≥ 0.7 were grouped as conserved, whereas those with lower scores were treated as non-conserved. (e) Classification of 3OST-1 *N*-glycosylation patterns based on sites number and positional configuration. In the pattern nomenclature, “x*N*Gly-y” denotes x predicted *N*-glycosylation sites, and y is the positional-configuration category based on the major recurrent arrangements of these sites in the multiple sequence alignment. Entries without a y index represent x groups in which only one major positional configuration was identified and no further subdivision was required. Brown boxes indicate highly conserved glycosylation sites, purple boxes indicate non-conserved glycosylation sites, and dark gray boxes indicate sporadic glycosylation-site positions.

Current strategies to address this issue have clear limitations that preclude their industrial translation. Reconstitution of *N*-glycosylation pathways in *E. coli* requires complex multistep engineering, and site-specific modification often remains incomplete (10–12). Direct removal of *N*-glycosylation sequons frequently disrupts folding and activity. Although advanced protein engineering strategies (including directed evolution, computational stability design, and fusion tag-assisted folding) have improved the prokaryotic expression of some eukaryotic proteins (13–15), these methods do not directly address their structural dependence on *N*-glycosylation. To date, there remains a critical lack of rational, structure-aware engineering strategies that can simultaneously overcome this glycosylation-induced expression bottleneck and preserve catalytic function to varying degrees while improving production-related performance.

For glycosaminoglycan (GAG) bioactive polysaccharides, chemoenzymatic biosynthesis has emerged as the dominant approach for their industrial-scale manufacturing(16–18). In this process, eukaryotic sulfotransferases are key rate-limiting enzymes because they govern both synthetic efficiency and product bioactivity through site-specific sulfation (19, 20). When expressed in *E. coli*, wild-type sulfotransferases often show limited soluble expression, poor functional stability, or reduced specific activity relative to eukaryotically expressed counterparts (21, 22). These limitations directly restrict the biological potency of biosynthetic GAG products and severely hinder their scalable industrial translation. Previous protein engineering efforts on the sulfotransferase family, including heparan sulfate *N*-deacetylase/*N*-sulfotransferase(22) (HS-NST), heparan sulfate 2-*O*-sulfotransferase(23) (HS-2OST), heparan sulfate 6-*O*-sulfotransferase(24) (HS-6OST), chondroitin sulfate 4-*O*-sulfotransferase(25) (CS-4OST), and chondroitin sulfate 6-*O*-sulfotransferase(26) (CS-6OST), mainly used conventional optimization, with limited targeted design for glycosylation-dependent constraints. Thus, this leaves an unmet need for targeted engineering strategies to overcome the glycosylation-dependent limitations on prokaryotic expression and residual activity stability for this enzyme family.

Inspired by the logic of ancestral state inference(27), we herein developed an evolution-guided optimization of *N*-glycosylation (EGO-*N*Gly) strategy, with heparin 3-*O*-sulfotransferase 1 (3OST-1) employed as the model target enzyme. 3OST-1 catalyzes the rate-limiting terminal 3-*O*-sulfation step in the chemoenzymatic biosynthesis of heparin, and is required for forming the antithrombin III-binding anticoagulant motif (28–30). However, wild-type 3OST-1 shows poor soluble expression and limited residual activity stability in prokaryotic hosts (31). EGO-*N*Gly strategy selectively reverts evolutionarily non-conserved *N*-glycosylation sequons to the naturally non-glycosylated residues conserved across early-divergent homologs, aiming to reduce glycan dependence and improve compatibility with *E. coli* expression. EGO-*N*Gly’s applicability was validated across 3OST-1 and other sulfotransferase family members(32, 33). Collectively, this work establishes EGO-*N*Gly as an evolution-guided engineering framework for improving the prokaryotic expression of selected glycosylation-dependent animal sulfotransferases.

## Results

### Evolutionary trajectory analysis of *N*-glycosylation sites in 3OST-1

Since *N*-glycosylation is a critical bottleneck in the heterologous expression of 3OST-1 in prokaryotic hosts, we analyzed *N*-glycosylation sequons (34) and systematically examined their evolutionary trajectory and sequence conservation (Figure 1b). The predicted *N*-glycosylation sites of 3OST-1 exhibit a clear phylogeny-associated conservation pattern(35), suggesting that different sites may be subject to distinct evolutionary constraints. Across the vertebrate orthologs analyzed, the number of predicted *N*-glycosylation sites generally increased from bony fish to amphibians and reptiles, and then decreased and became more stable in mammals, based on cross-species homolog mining, phylogenetic analysis (Figure S1, S2), and prediction of *N*-glycosylation sites (Figure 1c). In light of previous studies on protein thermodynamic evolution(36) and the stabilizing effects of glycan chains(37, 38), the enrichment of non-conserved glycosylation sites in some poikilothermic vertebrate lineages may reflect a possible auxiliary stabilization mechanism under fluctuating environmental conditions (39, 40). The establishment of endothermy in mammals and subfunctionalization of the 3OST family(41, 42) may have contributed to the reduced retention of these non-conserved sites in mammals(35, 43). However, these interpretations remain hypothetical, and additional comparative analyses will be required to distinguish the relative contributions of thermal adaptation, phylogenetic history, structural constraint, and functional diversification.

Phylogenetic mapping of the ortholog alignment identified seven recurrent candidate sequon positions, designated S1–S7 (Figure 1c). These alignment-defined positions represent candidate sites across the ortholog set and are not necessarily present in every ortholog. We calculated a site-specific conservation score as the frequency of the canonical N-X-S/T sequon (X ≠ Pro) at each aligned position. These *N*-glycosylation sites can be divided into two groups: highly conserved sites and non-conserved sites (Figure 1d, e). Using 0.7 as an operational working threshold for this dataset, sites with scores ≥ 0.7 were grouped as conserved, whereas those with scores < 0.7 were treated as non-conserved for the purpose of candidate stratification. On the basis of structural information and evolutionary evidence, we therefore proposed a glycosylation-site functional divergence model (Figure 2a): highly conserved sites are more likely to support core structural stability and catalytic function, whereas evolutionarily acquired non-conserved sites are more likely to contribute to eukaryote-specific folding and processing (44, 45). This interpretation is consistent with previous studies of chondroitin 4-*O*-sulfotransferase 1 (C4ST-1) and chondroitin 6-*O*-sulfotransferase 1(C6ST-1), in which conserved *N*-glycosylation sites were required for the formation of active and stable enzymes and localization to the Golgi apparatus, whereas disruption of these sites caused pronounced functional defects(46, 47). Accordingly, non-conserved sites are more suitable targets to improve expression in prokaryotic hosts compared with highly conserved sites(48, 49)

**Figure 2.**
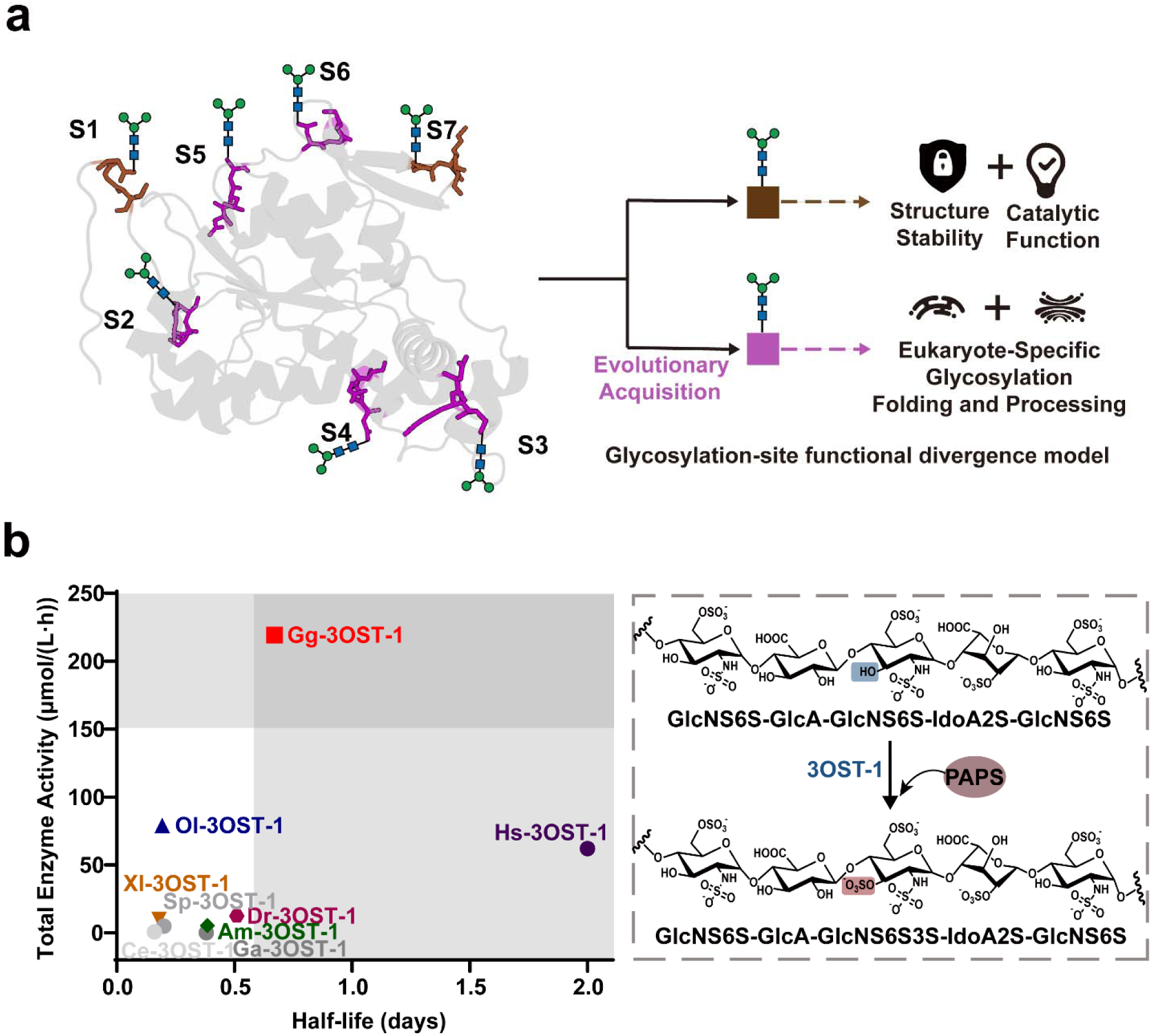
Glycosylation-site functional divergence model and comparative biochemical characterization of 3OST-1 orthologs. (a) Structural mapping of the seven *N*-glycosylation sites (S1-S7) and schematic model of glycosylation-site functional divergence. Conserved glycosylation sites are associated with structural stability and catalytic function, whereas evolutionarily acquired non-conserved sites are proposed to mainly facilitate eukaryote-specific folding and processing. (b) Quadrant plot comparing total enzyme activity and residual activity half-life at 37 °C of representative 3OST-1 orthologs. Total enzyme activity is defined as the overall catalytic output recovered from 1 L of culture, calculated as purified protein yield multiplied by specific activity, and expressed as μmol/(L·h). Residual activity half-life at 37 °C was calculated by fitting normalized residual activity decay curves to a first-order deactivation model. Each point represents the mean value from three independent biological experiments. Where shown, error bars indicate SD. Gg-3OST-1 showed the most favorable overall balance between activity output and stability among the tested orthologs and was therefore selected as the engineering scaffold. The inset shows the 3OST-1-catalyzed transfer of a sulfo group from 3’-phosphoadenosine-5’-phosphosulfate (PAPS) to the 3-OH position of a specific GlcNS6S residue in the substrate. Abbreviations: Am, *Apis mellifera*; Ce, *Caenorhabditis elegans*; Dr, *Danio rerio*; Ga, *Gasterosteus aculeatus aculeatus*; Gg, *Gallus gallus*; Hs, *Homo sapiens*; Ol, *Oryzias latipes*; Sp, *Strongylocentrotus purpuratus*; Xl, *Xenopus laevis*.

To compare ortholog performance under prokaryotic expression conditions, we evaluated three complementary metrics: specific activity, total enzyme activity, and residual activity stability. All proteins were purified and analyzed as MBP fusion constructs, and the MBP tag was retained for all biochemical and biophysical measurements. Here, specific activity refers to the specific activity of the purified enzyme, expressed as μmol product formed per mg enzyme per hour under the standard assay conditions (Figure S3). Purified protein yield refers to the amount of purified recombinant protein recovered from 1 L of bacterial culture and is expressed as mg/L. Total enzyme activity refers to the overall catalytic output recovered from 1 L of culture and was calculated as the product of purified protein yield and specific activity, expressed as μmol/(L·h). Residual activity stability was assessed by monitoring residual activity after separate pre-incubation of the purified proteins at 37 °C under the specified buffer conditions. Residual activity half-life at 37 °C was defined as the time required for residual activity to decrease to 50% of its initial value and was calculated from the fitted first-order deactivation rate constant derived from normalized residual activity decay curves. We therefore compared representative 3OST-1 orthologs using total enzyme activity and residual activity half-life, which reflects the persistence of catalytic function after 37 °C pre-incubation under the specified conditions (Figure 2b). Gg-3OST-1 (*Gallus gallus*) was selected as the primary engineering scaffold because it exhibited the most favorable overall performance among tested orthologs, including superior soluble expression, purified protein yield, and specific activity (Figure S3-S5), together with the highest total enzyme activity and a relatively long residual activity half-life (Figure 2b). This selection was based on overall biochemical performance rather than on the number of predicted *N*-glycosylation sequons alone. Indeed, naturally occurring orthologs containing only two predicted sequons, including Ce-3OST-1, Dr-3OST-1, and Sp-3OST-1, showed relatively low total enzyme activity and poor residual activity stability in *E. coli*, despite acceptable protein expression and solubility (Figures S4–S5). These observations indicate that reducing the number of predicted sequons is not sufficient to improve enzyme performance in an aglycosylated host.

### Implementation of the EGO-*N*Gly strategy synergistically enhances total enzyme activity and residual activity stability

Accordingly, evolution-guided optimizing of non-conserved *N*-glycosylation sites (EGO-*N*Gly) was designed as a selective remodeling strategy rather than an indiscriminate sequon-minimization strategy. It targets evolutionarily variable, non-conserved *N*-glycosylation sequon in a sequence background. To implement EGO-*N*Gly, we targeted non-conserved *N*-glycosylation sequons in Gg-3OST-1 and constructed a panel of combinatorial variants. Structural analysis showed that Gg-3OST-1 contains five predicted *N*-glycosylation sequons, including two conserved sites and three non-conserved sites, and its glycosylation-site profile belongs to the 5*N*Gly-1 cluster (Figure S6), where “5*N*Gly” indicates five predicted sequons and “-1” denotes a specific positional pattern of these sites. Guided by the EGO-*N*Gly strategy, we analyzed low-glycosylation orthologs in the 2*N*Gly-1 cluster and identified the corresponding highly conserved non-glycosylated residues (Figure S7). We then reverted the three non-conserved *N*-glycosylation sequons in Gg-3OST-1 to these residues (Figure 3a). Single, double, and triple variants were generated to evaluate combinatorial effects. Targeted reversion of non-conserved *N*-glycosylation sites progressively increased total enzyme activity and improved residual activity stability, with the triple variant showing the best overall performance. Specifically, Gg-3OST-M7 exhibited a 1.6-fold increase in total enzyme activity to 264 μmol/(L·h) (Figure 3b-d), mainly due to improved purified protein yield, while retaining WT-level specific activity. Its residual activity half-life at 37 °C increased 4.5-fold relative to wild-type (WT) (Figure 3e and Figure S8). The stability effects were non-additive. Although one single-site variant showed reduced stability, this defect was reversed in combined variants, suggesting epistatic interactions among the targeted sites and indicating that these positions jointly contribute to a coupled folding-stability network in the aglycosylated *E. coli* expression system(50). This pattern further suggests that simultaneous repair of multiple non-conserved sequon sites within a cluster may better offset host incompatibility than isolated single-site changes(51, 52).

**Figure 3.**
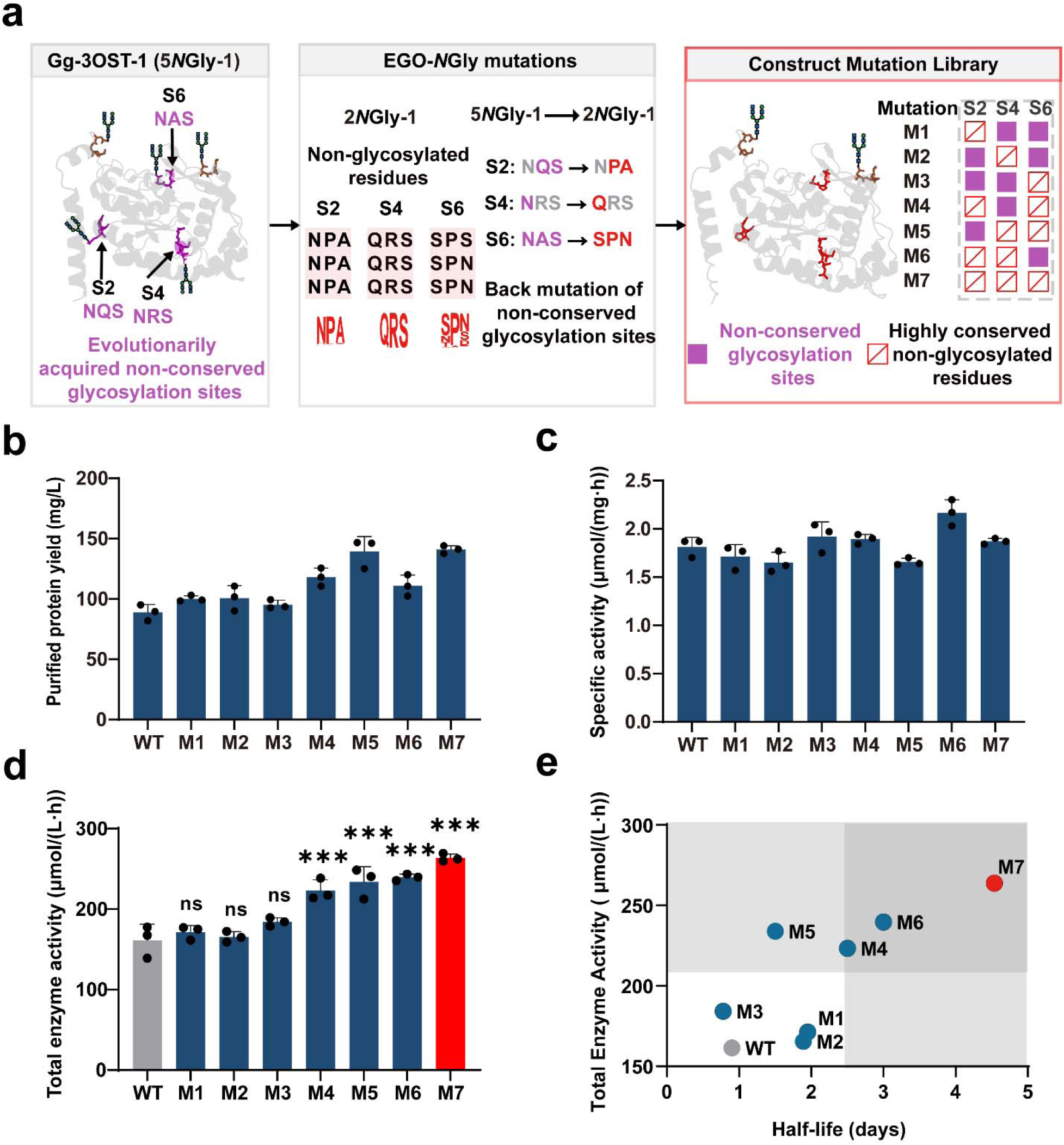
EGO-*N*Gly improves the total enzyme activity and residual activity stability of Gg-3OST-1. (a) Implementation of the EGO-*N*Gly strategy in Gg-3OST-1. Three evolutionarily acquired non-conserved *N*-glycosylation sites were reverted to the corresponding naturally non-glycosylated residues identified from early-diverging orthologs: S2, NQS→NPA; S4, NRS→QRS; and S6, NAS→SPN. The mutational states of S2, S4, and S6 in each variant are shown in the matrix. (b) Purified protein yield of wild-type Gg-3OST-1 (WT) and seven variants (M1–M7). (c) Specific activity of WT and the seven variants. (d) Total enzyme activity of WT and seven glycosylation-site mutations (M1–M7). Data are presented as mean values from three independent biological experiments (n = 3); upper error bars indicate SD. Individual data points are shown. Statistical significance was determined by one-way ANOVA followed by Tukey’s post hoc test (ns, not significant; *P < 0.05; **P < 0.01; ***P < 0.001; ****P < 0.0001). (e) Quadrant plot of total enzyme activity versus residual activity half-life at 37 °C for WT and corresponding variants. Each point represents the mean value from three independent biological experiments (n = 3). Where shown, upper error bars indicate SD. Residual activity half-life was determined by fitting normalized residual activity decay curves to a first-order deactivation model. Gg, *Gallus gallus*.

In contrast, substitution of the two highly conserved *N*-glycosylation sequons asparagine residues with glutamine reduced both purified protein yield and specific activity, leading to decreases in total enzyme activity to 0.57 and 0.39 of the WT level for the S1 and S7 mutants, respectively (Figure S9). Notably, because all variants were characterized in an aglycosylated *E. coli* host, these comparisons reflect the effects of the amino acid substitutions in the absence of eukaryotic *N*-glycosylation. At the three non-conserved sequon-related sites, simple N-to-Q substitutions also failed to reproduce the improved phenotype of the evolution-guided design. In Gg-3OST-1, the EGO-*N*Gly and N-to-Q designs coincide at S4, with both introducing N178Q; nevertheless, the EGO-*N*Gly triple variant increased total enzyme activity to 1.6-fold of WT, whereas the corresponding N-to-Q triple mutant retained only 0.7-fold of WT activity (Figure S10). These results indicate that the beneficial phenotype of M7 is not reproduced by simple conservative substitution at the targeted sequon asparagine residues. To test whether the M7 phenotype could instead be explained by generic surface remodeling or proline-mediated stabilization, we applied the same evolution-guided substitution strategy to a matched set of evolution-variable surface residues outside N-X-S/T sequons. These non-sequon control substitutions did not reproduce the overall gains observed for M7 and were often deleterious in combination (Figure S11), indicating that the beneficial effect depends on the positional context of the targeted sequon-related sites rather than nonspecific proline substitution alone.

Finally, to assess whether the observed improvement depended on the optimized expression background, we tested M7 in BL21(DE3) without GroEL/GroES co-expression. Although absolute production performance was substantially lower in this background, the relative advantage of M7 over WT was retained in both bacterial systems tested (Figure S12). Overall, these results show that EGO-*N*Gly enhances the total enzyme activity and residual activity stability of 3OST-1 in prokaryotic hosts via synergistic optimization of non-conserved *N*-glycosylation sites distal to the catalytic center, retains WT-level specific activity, and mitigating the common activity-stability trade-off observed in protein engineering(53, 54).

### Structural features underlying the enhanced stability of the EGO-*N*Gly variant

To elucidate the physicochemical mechanisms by which the EGO-*N*Gly strategy surmounts the folding threshold, we conducted thorough structural and dynamical analyses. Molecular dynamics (MD) simulations showed a significant reduction in the overall conformational flexibility of M7 (Figure S13, S14). Specifically, there were fewer fluctuations in three engineered regions of M7 compared with the WT according to root-mean-square fluctuation (RMSF) profiles (Figure 4a), which was consistent with the increased rigidity observed in biochemical experiments(55, 56). Globally, the surface electrostatic potential of M7 was redistributed (Figure 4b). The positively charged surface area decreased, and the overall potential decreased from 67 kT/e (WT) to 61 kT/e (M7), which likely reduces non-specific intermolecular interactions and helps maintain structural integrity during prokaryotic expression(57).

**Figure 4.**
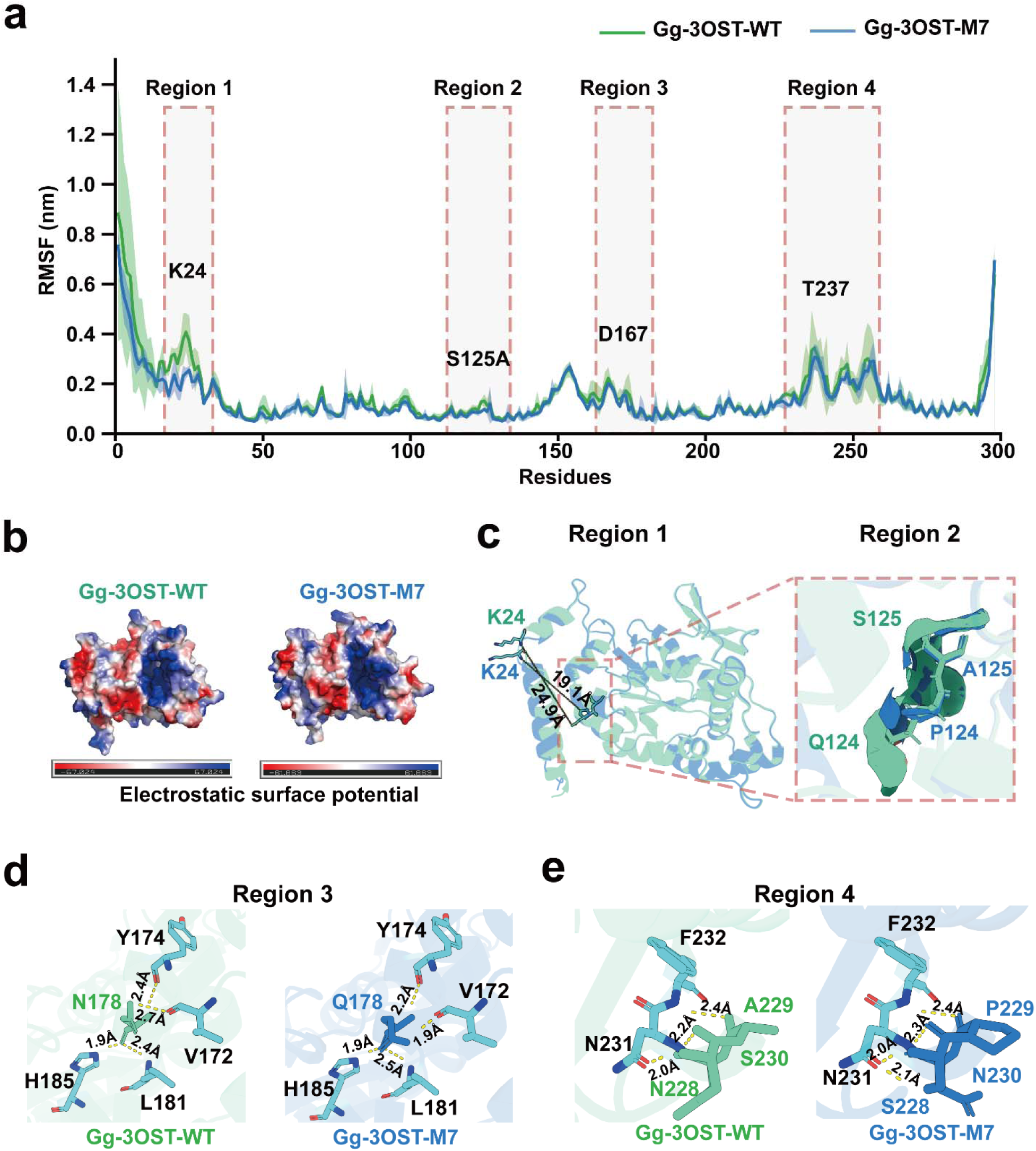
Structural basis for enhanced stability of the Gg-3OST-1 M7 variant. The WT and M7 are shown in green and blue, respectively. (a) Root-mean-square fluctuation (RMSF) profiles from molecular dynamics simulations comparing backbone flexibility in WT and M7, highlighting four regions with differential flexibility. RMSF profiles mean values from three independent 100-ns simulations (n = 3); shaded areas indicate SD. (b) Electrostatic surface representations of WT and M7 indicate a redistribution of surface electrostatic potential in the engineered variant. (c) Structural comparison of regions 1 and 2, showing conformational displacement in region 1 and local remodeling in region 2 associated with the Q124P/S125A substitutions. (d) Remodeling of the local hydrogen bonding network in region 3. Left, the WT N178-centered interaction pattern; right, the corresponding network in M7 after the N178Q substitution. (e) Reorganization of local interactions in region 4. Left, the WT N228–A229–S230 region; right, the corresponding region in M7 that carries N228S/A229P/S230N substitutions.

Further analysis of the four regions identified by RMSF showed that the mutations directly induced local network remodeling. In region 1 and region 2 (Q124/S125 loop), Q124P/S125A enhanced hydrophobic packing and restricted the backbone conformation, which reduces the distance between this loop and K24 from 24.9 Å to 19.1 Å (Figure 4c), and indicates that the flexibility is suppressed. In region 3, N178Q establishes a tighter hydrogen bond network with adjacent residues (including H185 and L181) (Figure 4d), which reinforces structural rigidity(58). In region 4, N228S/A229P/S230N reconfigures the hydrogen-bonding pattern to stabilize the adjacent α-helix (Figure 4e). S228 formed a novel 2.1 Å hydrogen bond and the pyrrolidine ring introduced by A229P restricted backbone flexibility(59, 60). Taken together, the structural and molecular dynamics analyses suggest that the combined substitutions in M7 reduce conformational fluctuations and remodel local intramolecular interaction networks. The increased hydrogen-bonding interactions and hydrophobic contacts observed in the M7 model are consistent with a more compact and conformationally constrained state. These changes may contribute to the improved stability and production-related performance of M7 in the aglycosylated expression system.

### Initial evaluation of the EGO-*N*Gly strategy across selected animal-derived sulfotransferases

To preliminarily assess whether EGO-*N*Gly can be applied beyond Gg-3OST-1, we tested a limited set of 3OST homologs from different vertebrate lineages and one member of the 6-*O*-sulfotransferase (6OST) family. Across these tested enzymes, the engineering outcome was strongly dependent on sequence context.

Cross-species testing of a limited number of 3OST orthologs showed that EGO-*N*Gly can improve total enzyme activity in some sequence backgrounds, but not uniformly across all tested enzymes. Following the workflow established above (Figure 5a), we systematically reverted evolutionarily acquired, non-conserved *N*-glycosylation sites in Xl-3OST-1(*Xenopus laevis*), Hs-3OST-1(*Homo sapiens*), Ol-3OST-1 (*Oryzias latipes*) and Am-3OST-1 (*Apis mellifera*) into their inferred natural non-glycosylated amino acids (Figure 5b). These reversions clearly produced background-dependent responses(61, 62): total enzyme activity increased by 2.9-fold in Hs-3OST-1, 1.6-fold in Xl-3OST-1 and, 2.3-fold in Am-3OST-1, whereas no significant change was observed for Ol-3OST-1 (Figure 5c,S15 and S16). In the responsive 3OST-1 orthologs, the increases in total activity were mainly associated with improved purified protein yield. Residual activity stability remained essentially unchanged for Hs-3OST-1, Xl-3OST-1, and Ol-3OST-1, whereas the Am-3OST-M1 showed a marked 5.8-fold increase in residual activity half-life. These results suggest that the main effects of EGO-*N*Gly are improvements in protein folding and expression in an aglycosylated prokaryotic environment. However, the magnitude and even the presence of this effect depend strongly on the local sequence and structural context of the target enzyme. The limited response of Ol-3OST-1 may arise from effects dependent on its sequence or structure(63). More broadly, these observations are consistent with possible epistatic effects, in which the outcome of a given glycosylation-site reversion depends on the overall sequence and structural background of each homolog. Notably, the present EGO-*N*Gly designs were restricted to reversions of evolutionarily acquired non-conserved *N*-glycosylation sites. Accordingly, some poorly expressing orthologs in our panel, including Dr-, Ce-, and Sp-3OST-1, already contained only the minimal two predicted *N*-glycosylation sequons and therefore were not prioritized for intervention in the present study. Thus, the current strategy is expected to be most applicable to orthologs that still retain removable non-conserved *N*-glycosylation sites, where it may confer beneficial effects.

**Figure 5.**
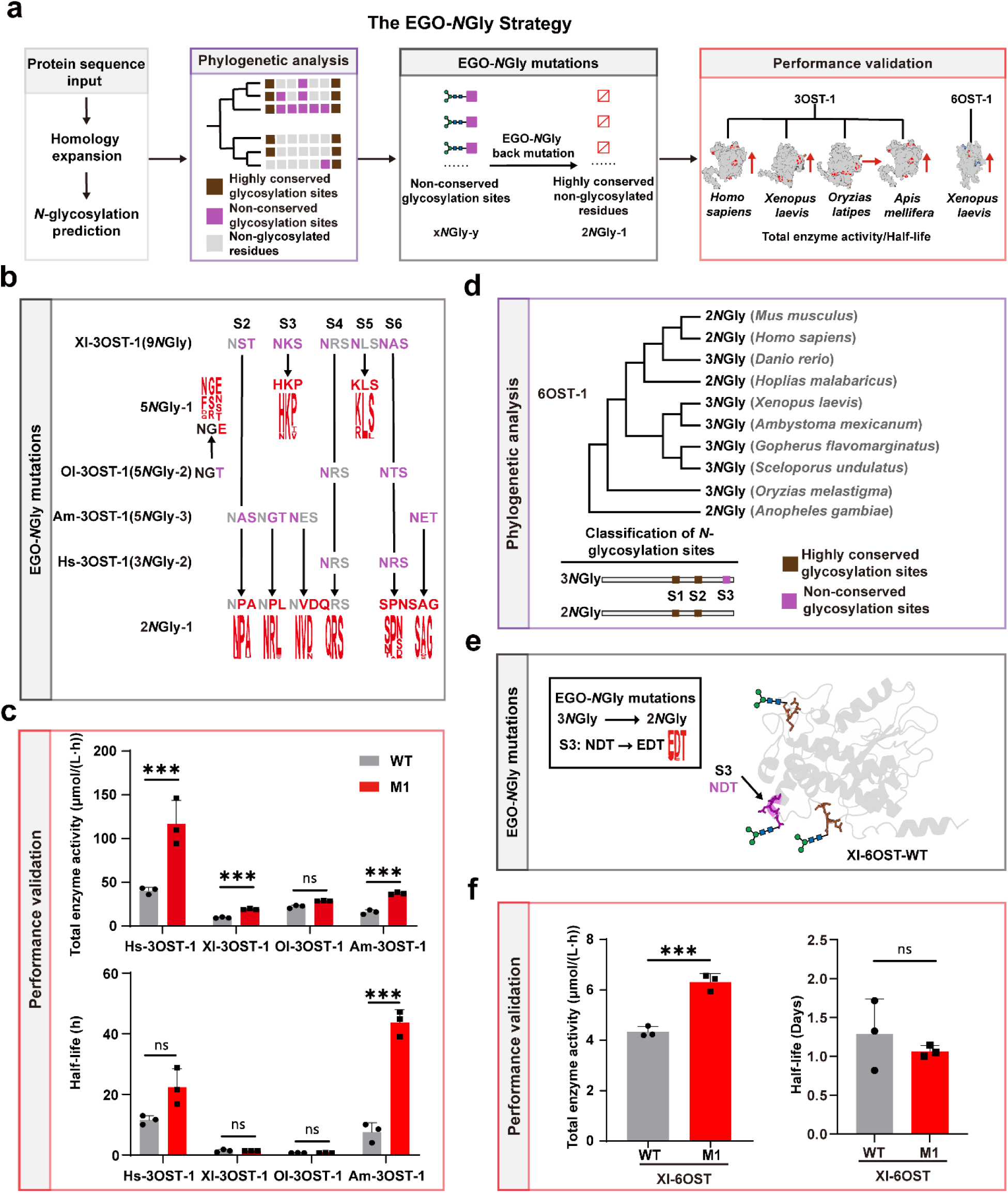
Transferability of EGO-*N*Gly across sulfotransferases, including 3OST and 6OST. (a) Workflow for EGO-*N*Gly-based remodeling of evolutionarily acquired non-conserved *N*-glycosylation sites, including protein sequence input, homolog collection, identification of non-conserved *N*-glycosylation sequons, phylogenetic analysis, back-mutation design, and experimental validation. (b) Sequence alignment and mutational design for representative 3OST-1 orthologs. (c) Total enzyme activity and residual activity half-life of wild-type (WT) and corresponding engineered mutant (M1) for representative 3OST-1 orthologs. Data are presented as mean values from three independent biological experiments (n = 3); upper error bars indicate SD. Individual data points are shown. Statistical significance was determined by two-tailed Student’s t test (ns, not significant; \**P* <0.05; \*\**P* <0.01, \*\*\**P* < 0.001). (d) Phylogenetic analysis and classification of *N*-glycosylation-site patterns in 6OST-1 orthologs, with conserved and non-conserved sites distinguished. (e) Structural mapping of *N*-glycosylation sites in Xl-6OST-1. The EGO-*N*Gly engineered S3 site (NDT→EDT) is highlighted. (f) Total enzyme activity and residual activity half-life of Xl-6OST-1 WT and the engineered mutant (M1) (n = 3). Data are presented as mean values from three independent biological experiments (n = 3); upper error bars indicate SD. Individual data points are shown. Statistical significance was determined by two-tailed Student’s t test (ns, not significant; \**P* <0.05; \*\**P* <0.01, \*\*\**P* < 0.001). 3OST, 3-*O*-sulfotransferase; 6OST, 6-*O*-sulfotransferase. Xl, *Xenopus laevis*; Ol, *Oryzias latipes*; Hs, *Homo sapiens*; Am, *Apis mellifera*.

We next tested whether EGO-*N*Gly could be extended across sulfotransferase families. We did not select 2OST or C5-epimerase because their *N*-glycosylation sites are highly conserved (Figure S17, S18), selected the 6OST family for validation because its evolutionary variation pattern is similar to that of the 3OST family (Figure 5d). Guided by EGO-*N*Gly, we used WebLogo to identify the consensus non-glycosylated residue from low-glycosylation orthologs in the 2*N*Gly-1 cluster (Figure S19). We then reverted the evolutionarily variable *N*-glycosylation site in Xl-6OST-1 to the corresponding non-glycosylated residue (Figure 5e). This single-point mutation increased the total enzyme activity by approximately 1.5-fold without measurable alterations in stability (Figure 5f and Figure S20), mainly through increased specific activity. Collectively, these results support the transferability of EGO-*N*Gly across related sulfotransferase families, and emphasize that its effect remains strongly dependent on the sequence background.

### EGO-*N*Gly Exhibits Orthogonal Compatibility with Established Protein Engineering Strategies

To test whether EGO-*N*Gly is compatible with other engineering strategies, we coupled it with the machine learning-driven PROSS(64) workflow. Structure-guided PROSS design based on Gg-3OST-WT generated seven candidate mutations for experimental evaluation (Figure 6a and Figure S21, S22). Among the EGO-*N*Gly mutations characterized above, the overall performance of M7 was comparable to representative PROSS variants, although it only required six targeted reversion mutations and retained over 95% sequence identity compared with the WT. This result supports the efficiency of EGO-*N*Gly as a minimal-perturbation strategy to improve host adaptation. Therefore, we combined M7 with D5, the PROSS variant showing the best balance between activity and stability, or with D7, the most stable PROSS variant, to generate SM1 and SM2, respectively (Figure 6a). D5 exhibited a residual activity half-life of 7 d, and D7 exhibited 9 d.

**Figure 6.**
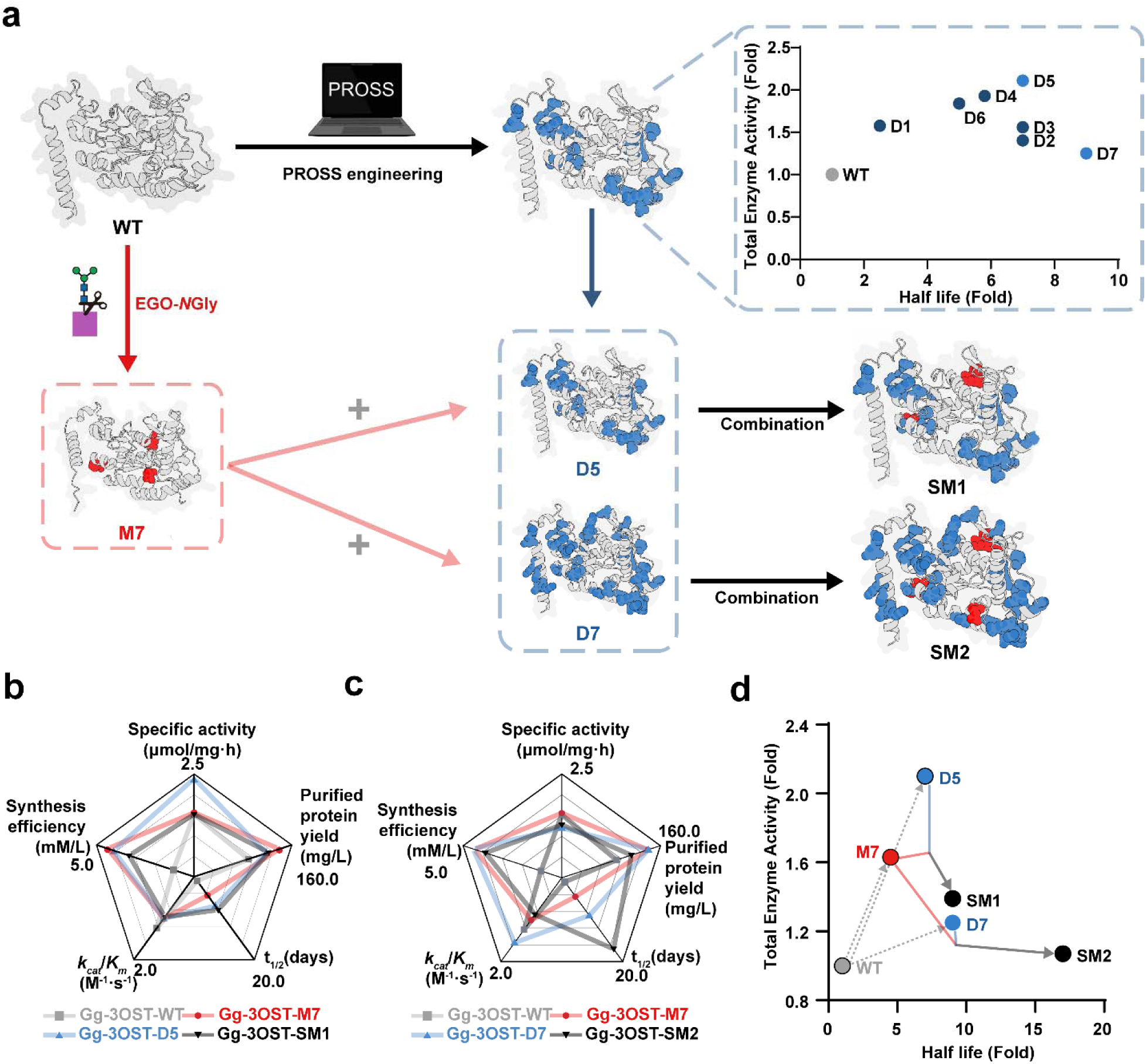
Orthogonal integration of EGO-*N*Gly and PROSS enhances the robustness of Gg-3OST-1. (a) Design workflow for combinatorial variants. Starting from wild-type Gg-3OST-1 (WT), the EGO-*N*Gly-derived variant M7 and PROSS-derived variants D5 and D7 were generated and combined to yield stacked variants: SM1 (M7 + D5) and SM2 (M7 + D7). The right panel shows a quadrant plot of relative changes in total enzyme activity and residual activity half-life for the initial PROSS variants (D1–D7) relative to WT. Values are shown as fold changes relative to WT and were calculated from mean measurements. (b, c) Radar plots summarizing the multidimensional performance metrics of SM1 (b) and SM2 (c), together with their parental variants and WT. Parameters include specific activity, product titer/synthetic efficiency, soluble protein expression, catalytic efficiency (*k*_cat_/*K*_m_), and residual activity half-life at 37 °C. Data are presented as mean values from three independent biological experiments (n = 3) for comparative visualization. (d) Scatter plot summarizing the trade-off between total enzyme activity and residual activity half-life across all engineered variants. Fold changes relative to WT were calculated from mean values obtained in three independent biological experiments and are shown for comparative visualization.

Both SM1 and SM2 showed improved overall biochemical performance relative to the WT, but their outcomes also revealed clear trade-offs among catalytic activity, functional expression, and robustness. SM1 exhibited a residual activity half-life of 8 d, exceeding those of M7 and D5, and achieved a 3-*O*-sulfated hexasaccharide product yield of 3.330 mmol/L culture, compared with 1.058 mmol/L for WT (Figure 6b, 6c and Figure S23). Because product yield integrates the effects of purified protein yield, specific activity, and enzyme persistence during the 18-h reaction, the improved yield of SM1 likely reflects the combined enhancement of these properties. Although SM1 showed higher functional expression than D5, its lower specific activity suggests that a modest catalytic cost is associated with the combined modifications, possibly owing to increased structural rigidity(54, 65). Thus, the gain in production-oriented performance in SM1 is accompanied by a measurable activity trade-off rather than a purely additive improvement. In parallel, SM2 exhibited a residual activity half-life of over 17 d, indicating substantially improved long-term robustness. Although the reduced specific activity of D7 was not fully recovered in SM2, its total enzyme activity reached 1.1-fold that of WT, together with a 3-*O*-sulfated hexasaccharide product yield of 3.869 mmol/L culture (Figure 6d). Collectively, these results show that product yield is determined by the combined contributions of purified protein yield, specific activity, and enzyme stability. Notably, all engineered variants retained catalytic efficiencies (*k*_cat_/*K*_m_) broadly similar to that of the WT within the uncertainty of the kinetic measurements (Table S1). Although modest numerical differences were observed among variants, their uncertainty ranges substantially overlapped with that of the WT, indicating no clear evidence of major catalytic impairment following either the EGO-*N*Gly reversions or the PROSS substitutions.

Taken together, these results suggest that EGO-*N*Gly and PROSS operate through distinct but complementary engineering criteria. EGO-*N*Gly primarily improves host adaptation by reverting evolutionarily variable non-conserved N-glycosylation sites. whereas PROSS enhances global thermodynamic stability through the distribution of stabilizing substitutions. Overall, the combination of EGO-*N*Gly and PROSS provides a tunable strategy for balancing functional expression, catalytic activity, and residual activity stability in the engineering of complex sulfotransferases.

## Discussion

This study establishes EGO-*N*Gly as an evolution-guided optimization strategy for improving the prokaryotic expression and residual activity stability of selected glycosylation-dependent sulfotransferases. In Gg-3OST-1, selective remodeling of non-conserved sequon increased total enzyme activity and substantially prolonged residual activity half-life, supporting the utility of this strategy for adapting animal-derived sulfotransferases to aglycosylated bacterial hosts. Mechanistically, MD simulations and structural analyses suggested that improved phenotype is associated with surface rigidity and remodeled intramolecular interaction networks. Consistent with this interpretation, differential scanning fluorimetry showed that representative engineered variants exhibited increased or maintained melting temperatures relative to WT (Figure S24). The apparent Tm of M7 increased from 70.36 °C to 71.83 °C, providing modest orthogonal biophysical support consistent with improved conformational stabilization of the fusion construct. Because DSF was performed on MBP-fused proteins, the measured Tm values represent the apparent stability of the fusion constructs rather than the intrinsic thermal stability of isolated 3OST. Importantly, the present data do not directly demonstrate glycan-removal effects per se. All constructs were expressed and characterized in an aglycosylated *E. coli* host, so the observed phenotypes reflect the consequences of the introduced amino acid substitutions in a non-glycosylating expression environment. Accordingly, EGO-*N*Gly should be interpreted here as an evolution-guided engineering strategy focused on non-conserved sequon-related positions and host compatibility.

EGO-*N*Gly improved total enzyme activity in several tested 3OST homologs and in one tested 6OST enzyme, suggesting that its design outcome may be influenced by the evolutionary and structural context of glycosylation-site clusters. Within the limited set of enzymes examined here, combinatorial site designs with stronger evolutionary conservation were associated with larger optimization gains than some single-site alterations. Broader sampling across additional sulfotransferases will be required to determine the generality of this pattern. More broadly, the effect of sequon reversion is unlikely to be determined by the glycosylation sequon alone, because neighboring residues and local structural networks may also have co-evolved to accommodate folding, stability, or expression constraints. Accounting for such co-evolved local interactions represents an important direction for refining the EGO-*N*Gly framework in future work.

A key feature of EGO-*N*Gly is that it introduces a minimal-perturbation design dimension centered on glycosylation-site evolution and host compatibility, making it complementary to current computational protein engineering methods. It shares the evolutionary logic of ancestral-state inference(66); but does not require the reconstruction of a full-length ancestral protein. Therefore, it differs from global sequence-redesign methods such as PROSS and ProteinMPNN(13), as well as local optimization strategies focused on the active site such as FuncLib(67). This study showed that EGO-NGly can be orthogonally integrated with PROSS, generating SM1 and SM2 with distinct trade-offs among activity, expression, and stability. This precision likely contributes to improved stability and functional compatibility in prokaryotic expression systems.

EGO-*N*Gly is not intended to recreate eukaryotic glycosylation in microbial hosts, but to improve the manufacturability and functional utility of animal-derived glycoprotein enzymes in cost-effective prokaryotic systems. By introducing limited sequence changes while retaining high WT sequence identity and catalytic activity, this strategy adapts animal-derived sulfotransferases for efficient biocatalytic production in aglycosylated hosts, rather than preserving human-like glycoforms as in mammalian glycoengineering. Engineering highly conserved sequon-associated residues, which may play essential structural roles, remains an open challenge and will likely require combining sequon reversion with compensatory stability design.

Overall, this work establishes a framework for engineering selected glycosylation-dependent sulfotransferases for improved prokaryotic production and yields more suitable enzyme candidates for the chemoenzymatic synthesis of glycosaminoglycans and heparins. However, the current validation remains limited to a small number of orthologs within the 3OST-1 family, while broader validation across additional orthologs and enzyme families will be required to define the generality of this approach. Even with these limitations, the present study provides a practical starting point for adapting selected animal-derived enzymes to microbial production systems.

## Materials and Methods

### Bioinformatics analysis

*Homo sapiens* 3-*O*-sulfotransferase-1 (Hs-3OST-1; GenBank: AAH57803), Hs-3OST-1 was used to identify homologs by BLASTP, which were further analyzed for signal peptides, putative N-glycosylation sites, phylogenetic relationships, and sequence similarity networks to guide ortholog selection, glycosylation-site comparison, and structure-based interpretation. Details are included in SI Appendix, Materials and Methods.

### Stepwise EGO-*N*Gly workflow and criteria for candidate-site selection

EGO-*N*Gly was implemented as a sequence- and evolution-guided workflow to identify non-conserved *N*-glycosylation sequons and redesign them by back-mutation to conserved non-glycosylated residue states. Details are provided in SI Appendix, Materials and Methods.

### Plasmid construction and expression host

Coding sequences of selected 3OST-1 orthologs, Gg-3OST-1, and derived mutants were cloned into pMAL-c5X, co-transformed with pGro7, and heterologously expressed in *Escherichia coli* Origami B (DE3) under chaperone-assisted, low-temperature induction conditions. Details are included in SI Appendix, Materials and Methods.

### Protein purification

Recombinant MBP-tagged 3OST-1 proteins were purified from cell lysates by amylose affinity chromatography and analyzed by SDS/PAGE for purity and apparent molecular weight. Details are included in SI Appendix, Materials and Methods.

### HPLC-based assay for 3OST-1 activity and product quantification

3OST-1 activity was quantified using an HPLC-based assay that monitored conversion of a *p*NP-labeled 6mer-6S acceptor substrate to the corresponding 3-*O*-sulfated product under standardized reaction conditions. Details are provided in SI Appendix, Materials and Methods.

### Specific activity, purified protein yield, and total enzyme activity

Enzyme performance was assessed by specific activity, purified protein yield, and total enzyme activity, with the latter calculated as the product of purified protein yield and specific activity. Details are provided in SI Appendix, Materials and Methods.

### Determination of 3-*O*-sulfated hexasaccharide product yield

Culture-level product output was estimated from HPLC-determined substrate conversion in a standardized reaction together with the purified enzyme yield recovered from 1 L of bacterial culture. Details are provided in SI Appendix, Materials and Methods.

### Residual activity stability and half-life determination at 37 °C

Residual activity stability was evaluated by measuring residual activity after pre-incubation at 37 °C and fitting the activity decay profiles to a first-order deactivation model to obtain residual half-lives. Details are provided in SI Appendix, Materials and Methods.

### Design and construction of variants

Conserved and nonconserved glycosylation-related sites were identified by sequence alignment and logo-based analysis, and single, double, and triple mutants were generated by site-directed mutagenesis and evaluated for expression, activity, and stability. Details are included in SI Appendix, Materials and Methods.

### Molecular dynamics simulation

Molecular dynamics simulations were performed on 3OST-1 systems to assess structural stability and conformational behavior using standard explicit-solvent protocols and trajectory-based structural analyses. Details are included in SI Appendix, Materials and Methods.

### PROSS engineering of wild-type Gg-3OST-1

The AlphaFold2-predicted structure of Gg-3OST-1 was used for PROSS-based stability design, and selected variants with favorable evolutionary and energetic profiles were experimentally tested for activity and thermostability. Details are included in SI Appendix, Materials and Methods.

### Enzyme kinetic parameters

Kinetic parameters of wild-type and engineered enzymes were determined from initial-rate measurements across substrate gradients and fitted to the Michaelis-Menten model. Details are included in SI Appendix, Materials and Methods.

### Differential scanning fluorimetry

Thermal stability of wild-type and engineered proteins was evaluated by differential scanning fluorimetry using purified MBP-fusion proteins. Details are provided in SI Appendix, Materials and Methods.

## Author Contributions

J.-Z.S., C.-Y.L. and J.-Q.D. conceived and designed the entire research project. C.-Y.L., Y.L. and X.-Y.L. explored and developed the methodology. C.-Y.L., J.-Q.D., and Y.-H.Z. conducted the experiments. J.-Y.S., X.-P.G. and Y.-L.C. provided essential materials and proteins. J.-Z.S., C.-Y.L. and S.Z. drafted the manuscript. All authors contributed to the manuscript, critically reviewed its content, and approved the final version for submission.

The manuscript was written through contributions of all authors. All authors have given approval to the final version of the manuscript.

## Acknowledgments

This work was supported in part by the National Key R&D Program of China (grant No. 2023YFA0914300), the National Natural Science Foundation of China (grant No. 82404458), the China Postdoctoral Science Foundation (grant No. 2025M773586) and Tianjin Major Science and Technology Special Project (grant No. 25ZXWCSY00120).

## Notes

## Abbreviations

2OST: 2-*O*-sulfotransferase
3OST-1: 3-*O*-sulfotransferase 1
6OST: 6-*O*-sulfotransferase
ASTIV-6: arylsulfotransferase IV-6
C4ST-1: chondroitin 4-*O*-sulfotransferase 1
C6ST-1: chondroitin 6-*O*-sulfotransferase 1
GlcNS6S: 2-N,6-*O*-disulfo-D-glucosamine
IPTG: isopropyl β-D-1-thiogalactopyranoside
PAPS: 3’-phosphoadenosine-5’-phosphosulfate
PNPS: Potassium p-nitrophenol sulfonate
PROSS: Protein Repair One-Stop Service
SSN: sequence similarity network

## Supporting Information

